# PDE1 inhibition modulates Ca_v_1.2 channel to stimulate cardiomyocyte contraction

**DOI:** 10.1101/2020.11.05.368852

**Authors:** Grace K Muller, Joy Song, Vivek Jani, Yuejin Wu, Mark E Anderson, David A Kass

## Abstract

**Rationale:** Cyclic adenosine monophosphate (cAMP) activation of protein kinase A (PKA) stimulates excitation-contraction coupling, increasing cardiac contractility. This is clinically leveraged by beta-adrenergic stimulation (β-ARs) or phosphodiesterase-3 inhibition (PDE3i), though both approaches are limited by arrhythmia and chronic myocardial toxicity. Phosphodiesterase-1 inhibition (PDE1i) also augments cAMP and was recently shown in rabbit cardiomyocytes to augment contraction independent of β-AR stimulation or blockade, and with less intracellular calcium rise than β-ARs or PDE3i. Early testing of PDE1 inhibition in humans with neuro-degenerative disease and dilated heart failure has commenced. Yet, the molecular mechanisms for PDE1i inotropic effects remain largely unknown.

**Objective:** Define the mechanism(s) whereby PDE1i increases contractility.

**Methods and Results:** Primary guinea pig myocytes which express the cAMP-hydrolyzing PDE1C isoform found in larger mammals and humans were studied. The potent, selective PDE1i (ITI-214) did not alter cell shortening or Ca^2+^ transients under resting conditions whereas both increased with β-ARs or PDE3i. However, PDE1i enhanced shortening with less Ca^2+^ rise in a PKA-dependent manner when combined with low-dose adenylate cyclase stimulation (Forskolin). Unlike PDE3i, PDE1i did not augment β-AR responses. Whereas β-ARs reduced myofilament Ca^2+^ sensitivity and increased sarcoplasmic reticular Ca^2+^ content in conjunction with greater phosphorylation of troponin I, myosin binding protein C, and phospholamban, PDE1i did none of this. However, PDE1i increased Ca_v_1.2 channel conductance similar to PDE3i in a PKA-dependent manner. Myocyte shortening and peak Ca^2+^ transients were more sensitive to Ca_v_1.2 blockade with nitrendipine combined with PDE1i versus PDE3i. Lastly, PDE1i was found to be far less arrythmogenic than PDE3i.

**Conclusions:** PDE1i enhances contractility by a PKA-dependent increase in Ca_v_1.2 conductance without concomitant myofilament desensitization. The result is less rise in intracellular Ca^2+^ and arrhythmia compared to β-ARs and/or PDE3i. PDE1i could be a novel positive inotrope for failing hearts without the toxicities of β-ARs and PDE3i.

## INTRODUCTION

Heart failure with depressed systolic function is a leading cause of morbidity and mortality and currently affects tens of millions of patients worldwide with a rising prevalence ^1^. Current drug treatments focus on reducing volume overload with diuretics and blocking β-adrenergic receptor (β-AR) and angiotensin stimulation. Methods to increase contractility have historically mimicked β-AR agonism to increase cyclic adenosine monophosphate (cAMP), which activates protein kinase A, though new methods to enhance sarcomere function directly are under development^2^. At present the most widely used therapeutics are the β-AR agonist dobutamine and phosphodiesterase type 3 (PDE3) inhibitor milrinone, the latter enhancing cAMP signaling by suppressing its hydrolysis. The inotropic effects from either are less potent in failing hearts as β-adrenergic receptor signaling and adenylyl cyclase activity are downregulated^3^. Their influence is further curtailed in many patients by the therapeutic use of β-AR blockade ^4^. Importantly, both approaches raise intra-cellular calcium and are pro-arrhythmic, factors which have constrained their use to acute indications ^5–7^. Safe and effective alternatives amenable to chronic therapy are lacking.

PDE1 is a dual cyclic nucleotide phosphodiesterase that is highly expressed in the mammalian heart. It is unique among PDEs because it requires calcium/calmodulin for activation. PDE1 is expressed as one of three isoforms, with PDE1C being most prominently expressed in human and other larger mammalian hearts. This isoform exhibits balanced selectivity for cAMP and cGMP, whereas isoform 1A, which is primarily expressed in rat and mouse heart, favors cGMP hydrolysis ^8, 9^. In 2018, we first reported that a pan-isoform inhibitor of PDE1 (ITI-214) increases contractility and reduces vascular resistance in conscious dogs with normal or failing hearts, and in intact rabbits ^8^. Our study further revealed that such effects were independent of β-AR stimulation (β-ARs) or blockade and changes in heart rate. Moreover, PDE1 inhibition (PDE1i) regulated cAMP differently than β-ARs or PDE3 inhibition (PDE3i), increasing myocyte contraction but with less intracellular calcium rise. These results have since led to a Phase Ib-IIa placebo-controlled study of ITI-214 in humans with dilated heart failure (NCT03387215). Preliminary results show positive inotropic and vasodilator effects much as found in dogs and rabbits^10^.

The intracellular mechanisms whereby PDE1i augments contractility remain unknown. PKA activation by β-AR stimulation leads to phosphorylation of the regulatory protein Rad to increase Ca_v_1.2 (L-type calcium channel) conductance ^11, 12^. Concomitant PKA phosphorylation of phospholamban (PLN) disinhibits the sarcoplasmic reticulum (SR) Ca^2+^ ATPase (SERCA) to increase SR calcium uptake and calcium-induced calcium release ^13, 14^. PDE3A localizes to the SR where it regulates cAMP-PKA stimulation ^13, 15^, so its inhibition can further augment SERCA2a activity. Collectively, these changes increase peak intracellular calcium transients and hasten their decline, improving contraction and relaxation. At the sarcomere, PKA phosphorylates troponin I (TnI) that desensitizes myofilaments to calcium and enhances relaxation ^16^, and myosin binding protein C (MyBP-C) which accelerates activation kinetics and increases β-ARs mediated contraction ^17, 18^.

Given that calcium transients are less altered by PDE1i, we hypothesized that intracellular PKA modulation and its consequences differ between β-ARs or PDE3i vs PDE1i. The current study tested this using guinea pig myocytes that also express the PDE1C isoform as in rabbits and humans. We find that the primary impact of PDE1i is to increase Ca_v_1.2 conductance without impacting PLN, TnI, or MyBP-C phosphorylation and associated with this, no change in SR calcium load or myofilament calcium sensitivity. The result is enhanced inotropy with less rise in intracellular calcium concentration and pro-arrhythmia than is observed with β-ARs and/or PDE3i.

## METHODS

### Reagents

The following reagents were used in the study: PDE1i - ITI-214 (provided under research agreement with Intra-Cellular Therapies, Inc, NY), cilostamide, forskolin and rolipram (Tocris, Minneapolis, MN), DMSO, Rp-cAMPS, caffeine, nitrendipine, verapamil (Millipore Sigma, Burlington, MA), and Rp-8-CPT-cAMPS (Cayman Chemical, Ann Arbor, MI). The following antibodies were used: PDE1A (1:1000, Sc-50480, H-105, Santa Cruz Biotechnology, Dallas, TX), PDE1C (1:5000, Ab14602, Abcam, Cambridge, MA), GAPDH (1:2000, 5174, Cell-Signaling, Danvers, MA), phospho-Ser^24/25^ troponin I (1:1000, ThermoFisher Scientific) and total troponin I (1:1000, ThermoFisher Scientific), phospho-Ser^273^, -Ser^282^, -Ser^302^ and total MyBP-C (1:2000, 1:4000, 1:4000, 1:2000, respectively, gifts from Dr. Sakthivel Sadayappan, University of Cincinnati), phospho-Ser16 phospholamban (1:2000, Badrilla, Leeds, UK), total phospholamban (1:3000, ThermoFisher Scientific, Grand Island, NY). To obtain single band detection with PDE1C Ab, we used 0.1% KPL (SeraCare, Milford, MA) as the blocking buffer. For all other immunoblots, we used Odyssey Blocking Buffer (Li-Cor, Lincoln, NE) 1:1 in TBST.

### Animals

Guinea pigs (N=57, males, 350-500 g) were used in this study. All animal study procedures were performed in accordance with the Guide to the Care and Use of Laboratory Animals and approved by the Johns Hopkins University Institutional Animal Care and Use Committee.

### Myocyte cell isolation

The protocol used for guinea pig myocyte isolation is described in detail elsewhere ^19^. Briefly, guinea pigs were anesthetized with pentobarbital, and hearts rapidly removed via thoracotomy. The aorta was cannulated on a Langendorff apparatus fitted with a heating jacket circulating water at 37 °C, and retrograde-perfused for 5 minutes at 8 ml/min with Tyrode’s solution. The perfusate was then switched to Tyrode’s solution containing collagen type 2 (Worthington, Columbus, OH) and protease type 14 (Sigma-Aldrich, St. Louis, MO) for 7-9 minutes. The solution was switched to a modified Kraft-Bruhe (KB) buffer for 5 minutes, before hearts were minced and filtered (200 μm) to yield single cells. Myocytes were then rested in modified KB buffer ^20^ for one hour before being placed in supplemented M199 ACCIT medium ^21^. Cells were maintained at room temperature and studied over 7 hours.

### Sarcomere shortening and Ca^2+^ transients

Changes in cell sarcomere shortening and Ca^2+^ transients were measured using a customized IonOptix system previously described ^8^. All recordings were performed at 37° C, with pacing stimulation at 1 Hz. Cells were loaded using 1 μM Fura-2 for 15 minutes and then washed for at least 20 minutes. Baseline recordings were made in Tyrode’s buffer with 0.1% DMSO. Cells were subsequently stimulated with various agents as described in results. Three baseline parameters were considered for the two types of measurements (pre-stimulation, peak percent change after stimulation, and time to return to 50% baseline). Cells falling within 2 standard deviations of the mean value were included in group analysis. To determine the role of PKA in PDE1i or PDE3i mediated response, cells were pre-incubated with 100 μM Rp-8-CPT-cAMPS for 35-45 minutes ^22^ before being studied. To assess SR Ca^2+^ content, caffeine-induced SR Ca^2+^ release was performed as previously described ^23^. For this assay, stable responses to forskolin+ITI-214 or Iso, or DMSO in normal Tyrode’s were recorded for one minute prior to stopping pacing, and caffeine (10 mM) introduced via a needle placed adjacent to the cell. The peak Ca^2+^ transient was quantified.

### Myofilament force-pCa relationship in skinned myocytes

Myocytes were incubated in 0.3% Triton X-100 in isolation buffer with protease inhibitor (Sigma-Aldrich) and phosphatase inhibitor (PhosSTOP, Roche, Indianapolis, IN) for 20 minutes at 10°C as described earlier ^24^. After washing in isolation buffer, cells were attached with UV-activated adhesive (Norland Products, Cranbury, NJ) to a force transducer-length controller (Aurora Scientific, Aurora, Canada). Sarcomere length was adjusted to 2.1 *μ*m with micromanipulators (Siskiyou Corporation, Grants Pass, OR) as measured by digital 2D fast Fourier transform of images (IPX-VGA210, Imperx, Boca Raton, FL). Tension was equal to force divided by myocyte cross sectional area. Active tension-Ca^2+^ relationships were generated by varying Ca^2+^ concentration from 0 to 46.8 μmol/L. Tension -log[Ca^2+^] relations were fit to the Hill equation to obtain maximal tension (T_max_), Ca^2+^ sensitivity (EC_50_), and Hill coefficient^25^. Tension-pCa relationships were normalized to T_max._

### L-type Ca^2+^ current density recordings

Ca_v_1.2 current (I_Ca_) was measured with whole-cell patch clamp at 34±1 °C, as reported^26^. I_Ca_ was confirmed by sensitivity to nitrendipine (10 μM). Depolarizing voltage pulses (300 ms in duration) to potentials ranging −70 to 60 mV, in 10 mV steps, were applied from a holding potential of −80 mV. A pre-pulse to −40 mV of 50 ms was applied before each step to inactivate Na^+^ currents. An example voltage protocol is shown in Fig 4a. The pipette (intracellular) solution consisted of (in mM): 120 CsCl, 10 HEPES, 10 tetraethylammonium (TEA) chloride, 1.0 MgCl_2_, 1.0 NaGTP, 5.0 phosphocreatine, 3.0 CaCl_2_, 10 EGTA; pH was adjusted to 7.2 with 1.0 N CsOH. For PKA inhibition, Rp-cAMPS (100 μM) was included in the pipette. The bath (extracellular) solution consisted of (in mM): 137 NaCl, 10 HEPES, 10 glucose, 1.8 CaCl_2_, 0.5 MgCl_2_, and 25 CsCl, with pH adjusted to 7.4 with NaOH.

### Western blots

To probe for PLN phosphorylation, guinea pig myocytes were incubated with DMSO or various stimulation drugs all dissolved in Tyrode’s with 1.8 mM Ca^2+^ and rotated in a cell suspension for five minutes at room temperature. Cells were then quickly spun down, Tyrode’s replaced with lysis buffer (Lysis Buffer, Cell Signaling), and homogenized using beads and mechanical shearing at 30 Hz for 2 min (Retsch, Newtown, PA). The lysates were clarified using centrifugation (2000xg, 10 min) and assayed for protein content (BCA Assay, Thermo-Fisher). For sarcomeric proteins, cells were allowed to settle for three hours on laminin-coated 6-well plates. The M199 ACCIT media was replaced with identical media without bovine serum albumin (BSA) 30 minutes before the drug incubation. Cells were treated with DMSO or indicated drugs dissolved in the BSA-less media and incubated for 10 minutes at room temperature. Sarcomere fractions were isolated and quantitated as reported before ^24^. Proteins were separated on gel electrophoresis, followed by hybrid wet transfer onto nitrocellulose. 12% gels were used for phospholamban; 4-15% gels were used for PDE1A, PDE1C, GAPDH, troponin I and MyBP-C. Li-Cor Imager was used to scan and quantitate densitometry values.

### Statistical analysis

Raw data were collated using Excel and analyzed and graphed using Prism 8.0. Figures generally display all the individual data along with median and 25/75% in the form of violin plots, or individual data with mean +/− SEM as bar graphs. Other formats and statistics are noted along with statistical tests used in each of the figure legends.

## RESULTS

### Effects of PDE1-I on cAMP-stimulated contraction in guinea pig myocytes

PDE1 isoform expression has not been previously reported in guinea pig, so we first assessed and confirmed that PDE1A and PDE1C protein are both robustly expressed (Supplemental Fig 1). This pattern is similar to that reported in rabbit and human^8^. Figure 1A-C shows myocyte sarcomere shortening and Ca^2+^ transients at rest and upon exposure to the β-AR agonist isoproterenol (Iso, 1 nM), PDE3i cilostamide (Cil 1 μM), PDE4i rolipram (Rol, 10 μM), or PDE1i (ITI-214, 1 μM). Both Iso and Cil increased shortening and Ca^2+^ transients similarly. In contrast, neither Rol nor ITI-214 had any effect. The lack of PDE1i response in guinea pig myocytes is similar to data from rabbit^8^. The lack of PDE4i responses similar to results from larger mammalian hearts but while contrasting to mouse or rat for which PDE4 dominates EC coupling^27^.

**Fig 1.**
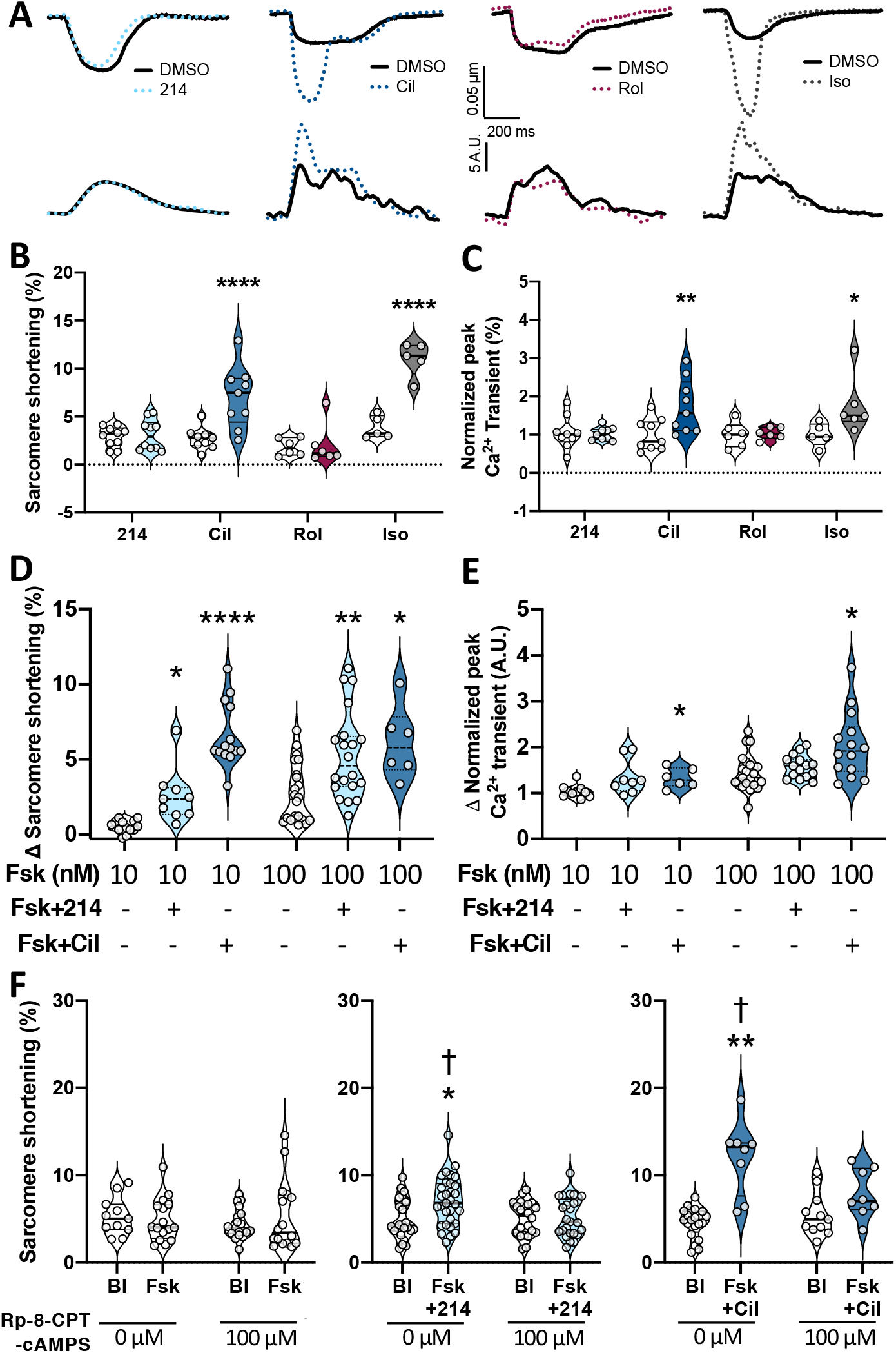
PDE1-I stimulates contraction at less Ca^2+^ rise than PDE3-I in guinea pig. Myocytes were treated with inhibitors targeting PDEs 1, 3, or 4: ITI-214 (214; 1 μM), cilostamide (Cil; 1 μM), and rolipram (Rol; 10 μM), respectively; or with isoproterenol (Iso; 1 nM). A) Representative traces showing changes in cell sarcomere shortening (upper traces), and Ca^2+^ transients (lower traces). Peak changes in B) sarcomere shortening and C) Ca^2+^ transient amplitude are plotted in a paired fashion against DMSO (open dot). *p<0.05, **p<0.01, ****p<0.0001 against respective DMSO; RM 2-way ANOVA with Sidak’s multiple comparison test. A separate group of cells were treated with forskolin (Fsk; at 10 or 100 nM), with or without additional 214 or Cil. The group averaged delta values for D) sarcomere shortening and E) Ca^2+^ transients from DMSO are plotted. *p<0.05, **p<0.01, ****p<0.0001 against respective DMSO; Kruskal-Wallis test. F) Change in sarcomere shortening was compared in the absence or presence of 100 μM Rp-8-CPT-cAMPS for cells treated with Fsk (10 nM), Fsk (10 nM) + 214 (1 μM), or Fsk (10 nM) + Cil (1 *μ*M). *p<0.01, **p<0.001 vs corresponding baseline (Bl), †p<0.05 vs corresponding drug condition in the presence of Rp-8-CPT-cAMPS; ordinary 2-way ANOVA with Sidak’s test.

Since inotropic effects from PDE1i require sufficient ambient cAMP that can in turn be modulated^28^, we next examined the response to PDE1i and PDE3i in cells pre-treated with the adenylate cyclase stimulator forskolin (Fsk). Based on dose response data (Supplemental Fig 2A), cells were exposed to either 10 or 100 nM Fsk alone or in combination with PDE1i or PDE3i (Figure 1D, 1E). PDE1i significantly augmented shortening but not Ca^2+^ transients whereas PDE3i increased both at either dose of Fsk pre-stimulation. To test whether inotropic enhancement by PDE1i requires PKA activation, we repeated this study using 10 nM Fsk pre-stimulation in the presence or absence of PKA inhibition (Rp-8-CPT-cAMPS, 100 μM). Consistent with prior data, PKA inhibition alone did not alter baseline sarcomere shortening^22^ but slowed relaxation kinetics (Supplemental Fig 2C). However, PKA inhibition blocked increased contraction (Figure 1F) and enhanced relaxation (Supplemental Fig 2D) from PDE1i and PDE3i.

Lastly, we examined the effect of PDE1i and PDE3i superimposed on β-AR stimulated shortening and Ca^2+^ transients. Both parameters were further augmented by PDE3i whereas PDE1i did not amplify the changes from β-ARs (Figure 2A, 2B). Taken together, these studies identify differences in myocyte function and calcium responses to PDE1i versus PDE3i, but similar dependence on PKA activation.

**Fig 2.**
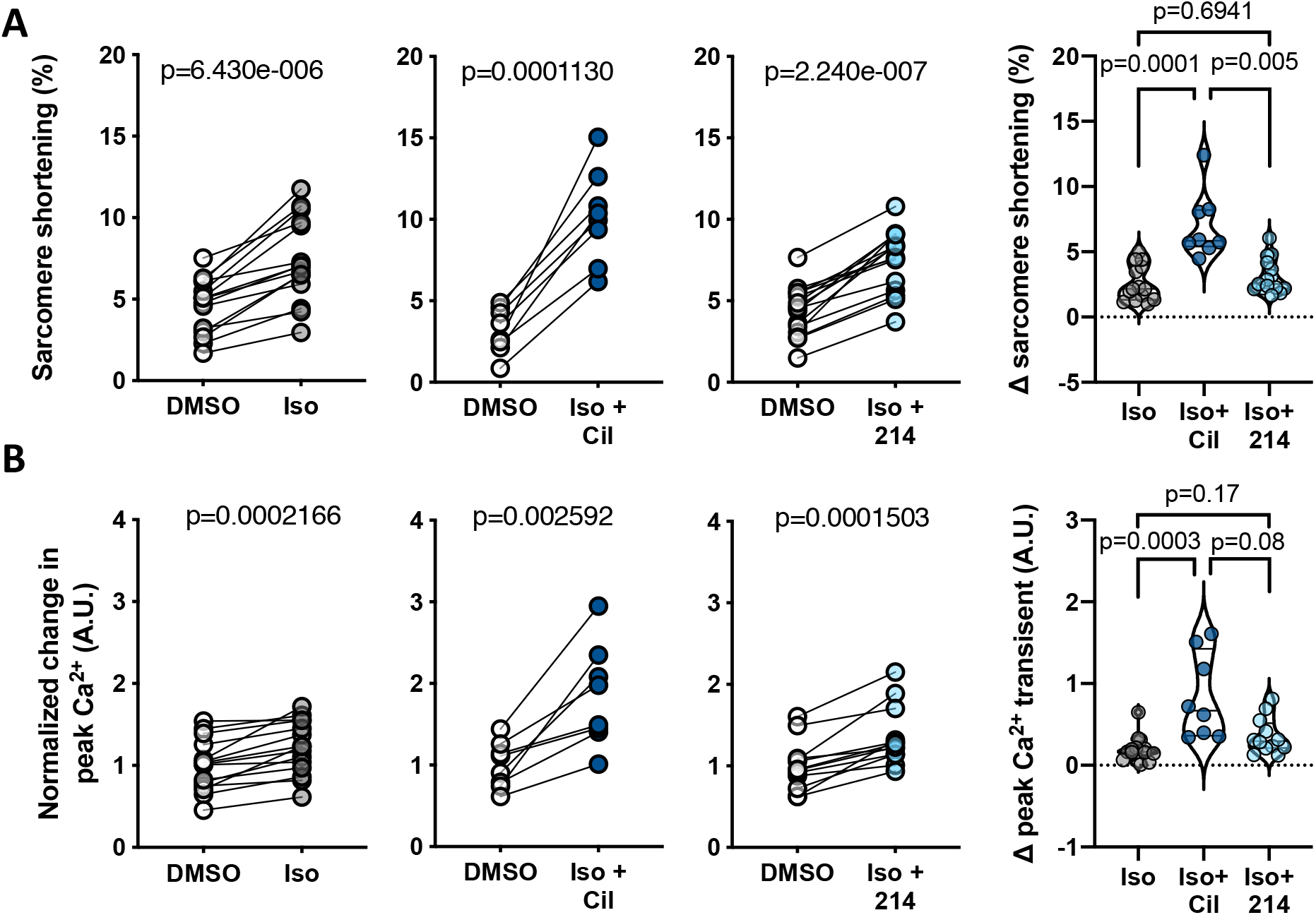
PDE1-I vs PDE3-I modulation of β-AR-stimulated signaling. Guinea pig myocytes were treated with sub-maximal isoproterenol (Iso; 0.025 nM) alone, or with Cil or 214. Changes in the peak A) sarcomere shortening and B) Ca^2+^ transients are plotted, with p-values indicating paired T-test results. The delta differences are plotted to the right. For the delta comparison, p values indicate the results of a Kruskal-Wallis test.

### Unlike β-AR stimulation, PDE1-I does not alter myofilament tension -Ca^2+^ relations

Beta-AR stimulation results in phosphorylation of TnI at Ser^23/24^, causing a rightward shift of the myocyte tension-Ca^2+^ dependence (myofilament desensitization) without changing maximal tension 29. This is characterized by greater Ca^2+^ required for 50% activation (EC_50_). We speculated that by contrast, PDE1i may not desensitize the myofilaments since it augments shortening at less Ca^2+^ elevation. To test this, myocytes incubated for 5 minutes with DMSO, Iso (50 nM), or Fsk (10 nM) + ITI-214 (1 μM) were placed in skinning solution with phosphatase inhibitors, and mounted on a force-length control apparatus to obtain normalized tension (force/cross sectional area) - Ca^2+^ relations. Stimulation with Iso shifted the curve rightward (desensitization) as expected, increasing EC_50_ from 2.42 to 4.61 (p=0.02). However, PDE1i had no significant impact on the relation and EC_50_ (Figure 3A, 3B). Maximal activated tension and Hill cooperativity were unaltered by either intervention (Table 1).

**Table 1.** Myofilament parameters in skinned guinea pig myocytes

|  | **DMSO** | **Iso** | **Fsk+214** |
| --- | --- | --- | --- |
| **Fmax** | 20 ± 1.9 | 23 ± 3.6 | 16 ± 2.4 |
| **EC<sub>50</sub>** | 2.4 ± 0.12 | 4.6 ± 0.69 * | 2.8 ± 0.57 |
| **Hill's coefficient** | 5.5 ± 1 | 3.3 ± 0.58 | 3.9 ± 0.69 |

\* p=0.02 vs DMSO; ordinary 1-way ANOVA with Dunnett's test

**Fig 3.**
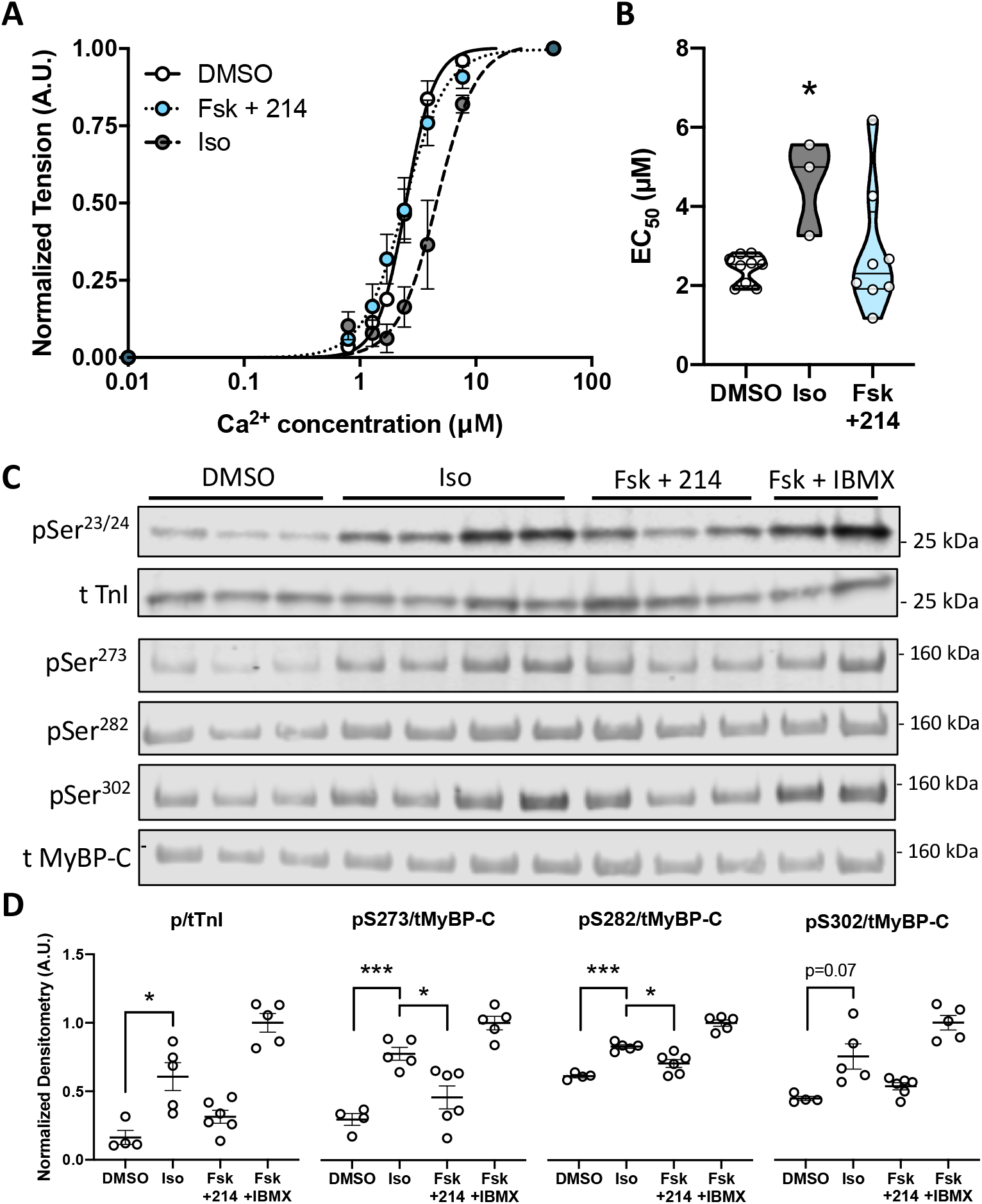
PDE1 inhibition does not alter the myofilament force-pCa relationship. Guinea pig myocytes were treated with DMSO, Fsk (10nM) + 214 (1 μM), or Iso (50 nM) before being skinned. A) A normalized curve showing the myofilament force-pCa relationship. B) The EC_50_ values, indicative of Ca^2+^ at 50% maximal activation, are plotted for these groups. C) Representative western blot of phosphorylated and total troponin I (TnI) at Ser^23/24^ or myosin binding protein-C (MyBP-C) at Ser^273^, Ser^282^ and Ser^302^ for guinea pig myocytes treated as indicated. The averaged group response, normalized to the saturating level of Fsk (25 μM) + IBMX (100 μM), is shown below. *p<0.05, **p<0.01; Brown-Forsythe and Welch ANOVA with Dunnett’s T3 multiple comparisons test.

**Fig 4.**
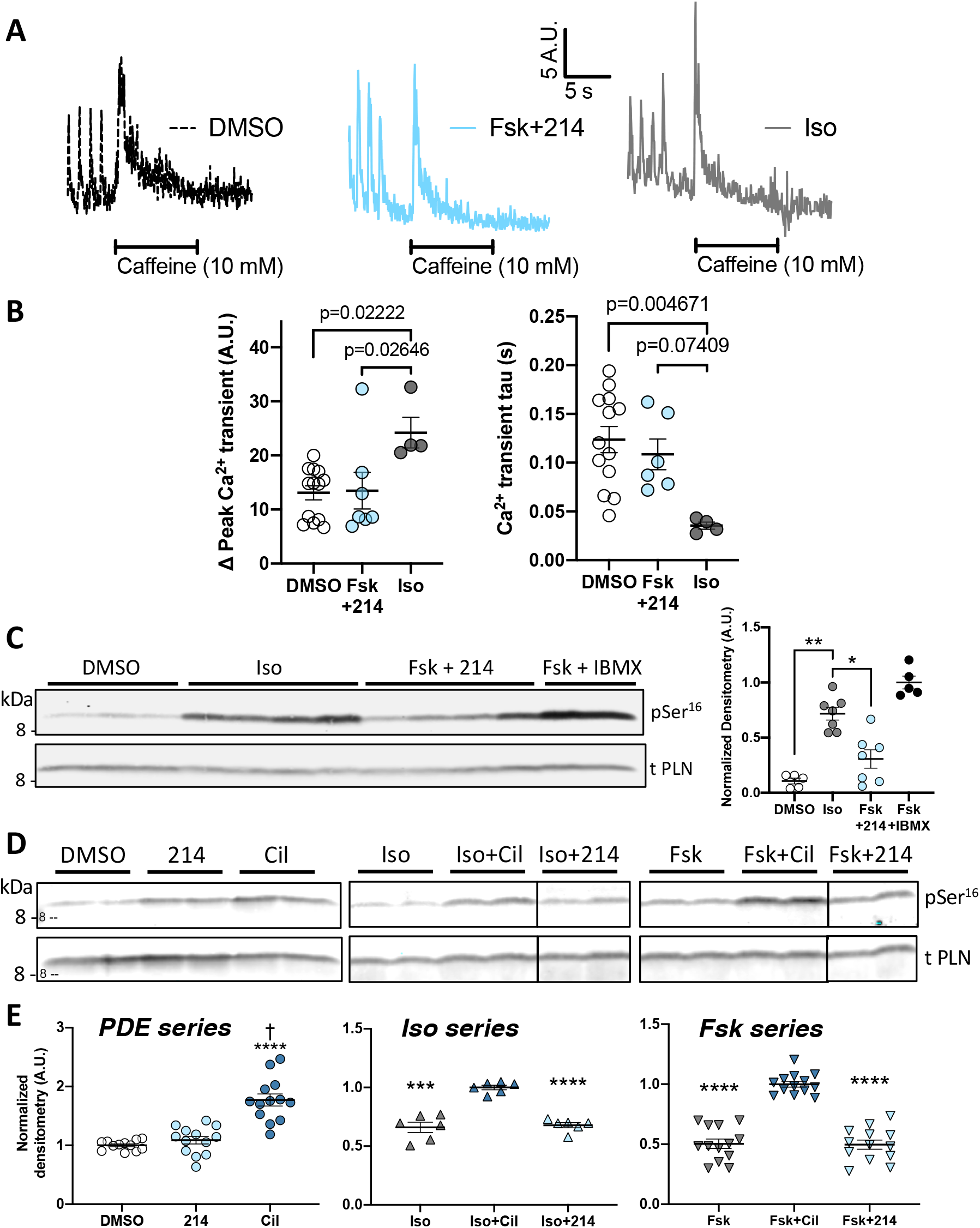
PDE1 inhibition does not increase the SR Ca^2+^ content, or the phospholamban (PLN) Ser^16^ phosphorylation level. Guinea pig myocytes treated with DMSO, Fsk (10nM) + 214 (1 μM) or Iso (1 nM) were treated with caffeine (10 mM). A) Representative Ca^2+^ traces are shown. B) Grouped average changes in the peak Ca^2+^ transients and relaxation constant tau are plotted, with p-values indicating Kruskal-Wallis test results. Representative western blot of phosphorylated (Ser^16^) and total PLN for myocytes treated with Fsk (10nM) + 214 (1 μM) in comparison to those treated with C) Iso at near maximal dose (50 nM) with quantiation to the right, or E) Iso at non-saturating dose (1 nM). E) Grouped average data are plotted. The level of phosphorylated/total PLN was normalized to that of Fsk+IBMX in D, or to that of DMSO, or Iso+cil or Fsk+cil in F, going from left to right. **** p<0.0001 vs DMSO, † p=0.0005 vs 214 in PDE series; *p<0.05, ** p<0.01, **** p<0.0001 vs Fsk+cil or Iso+cil in the rest; Kruskal-Wallis test.

The lack of altered tension-Ca^2+^ dependence suggested PKA phosphorylation of myofilament proteins, notably TnI and MyBP-C that occur with β-AR stimulation, would be lacking with PDE1i. Figure 3C and D show example immunoblots and summary data, using the same doses that induced similar physiological stimulation. The data are normalized to the maximal response obtained with Fsk+IBMX (100 μM). TnI phosphorylation with Iso was significantly greater than with Fsk+ITI-214. MyBP-C phosphorylation upon β-AR stimulation is important to enhancing contractility^30^. As PDE1i did not amplify pre-existing β-AR stimulation, we speculated this too did not occur. Iso phosphorylated MyBP-C Ser^273^, Ser^282^, and Ser^302^; however, this was not observed with Fsk+ITI-214.

### PDE1-I does not increase SR Ca^2+^ load or phospholamban phosphorylation

A major source of increased Ca^2+^ with β-AR is from enhanced SR release coupled to increased SR stores. Since PDE1i augmented contraction with less Ca^2+^ rise, we speculated these SR changes may not occur. Myocytes were treated with Fsk+ITI-214 or Iso in normal Tyrode’s and then exposed to caffeine (10 μM) to induce SR Ca^2+^ release. Fig 4A displays example Ca^2+^ response data for caffeine-induced SR Ca^2+^ storage analysis ^31^, and Figure 4B, summary data for the peak Ca^2+^ response reflecting SR Ca^2+^ load. Iso (1 nM) increased SR Ca^2+^ storage above vehicle control (DMSO), but Fsk+ITI-214 did not. Iso also shortened the Ca^2+^ decay constant (0.035 ± 0.0036 s) compared to DMSO (0.12 ± 0.013 s; p<0.0047, Kruskal-Wallis test), and this was unaltered by Fsk+214 (0.11 ± 0.016 s) (Figure 4B).

PKA-mediated PLN phosphorylation at Ser^16^ plays a major role in enhancing SR Ca^2+^ uptake; therefore, we further examined if Iso and Fsk+ITI-214 differed with respect to this post-translational modification. Myocytes exposed to Iso (50 nM) showed a rise in Ser^16^ phosphorylation that was not found with Fsk+ITI-214 (Fig 4C). This was further explored in cells treated with PDE1i or PDE3i alone, or in combination with non-saturating levels of Iso or Fsk (Figure 4D, 4E). In all conditions, PDE3i augmented PLN phosphorylation over the prior baseline significantly more than PDE1i.

### Ca_v_1.2 cannel current increases with both PDE1- and PDE3-I

The lack of myofilament or SR modifications by PDE1i led us to consider the opening of Ca_v_1.2 – the key first step in EC coupling. As recently revealed by the Marx laboratory ^11, 12^, PKA phosphorylates the binding protein Rad to dis-inhibit the channel and increase conductance. In larger mammalian cells such as those from dog, rabbit, and guinea pig, the relative role of Ca_v_1.2 versus SR Ca^2+^ is 1:2 and near 1:1 in failing hearts ^32, 33^. If PDE1i enhanced this current without inducing myofilament Ca^2+^ desensitization, it could augment contractility even without an SR component. We determined nitrendipine-sensitive I_CaL_ current density by voltage clamp protocol and average current density plotted versus depolarizing voltage (Fig 5A). Compared to DMSO, the I-V plot remained unaltered by Fsk or ITI-214 alone. However, combining PDE3i or PDE1i with low-dose Fsk (10 nM) resulted in a marked increase in current density (Fig 5B). Summary data for peak current (Fig 5C) confirmed similar enhancement with either PDE inhibitor. A maximal dose of nitrendipine (10 μM) blocked the current increase elicited by either of the two PDE inhibitors. To test if this effect was PKA dependent, studies were repeated with the PKA inhibitor Rp-cAMP (100 μM) in the recording electrode to dialyze the inhibitor into the cell. PKA inhibition blocked increased current from PDE1i and PDE3i +Fsk (Fig 5D).

**Fig 5.**
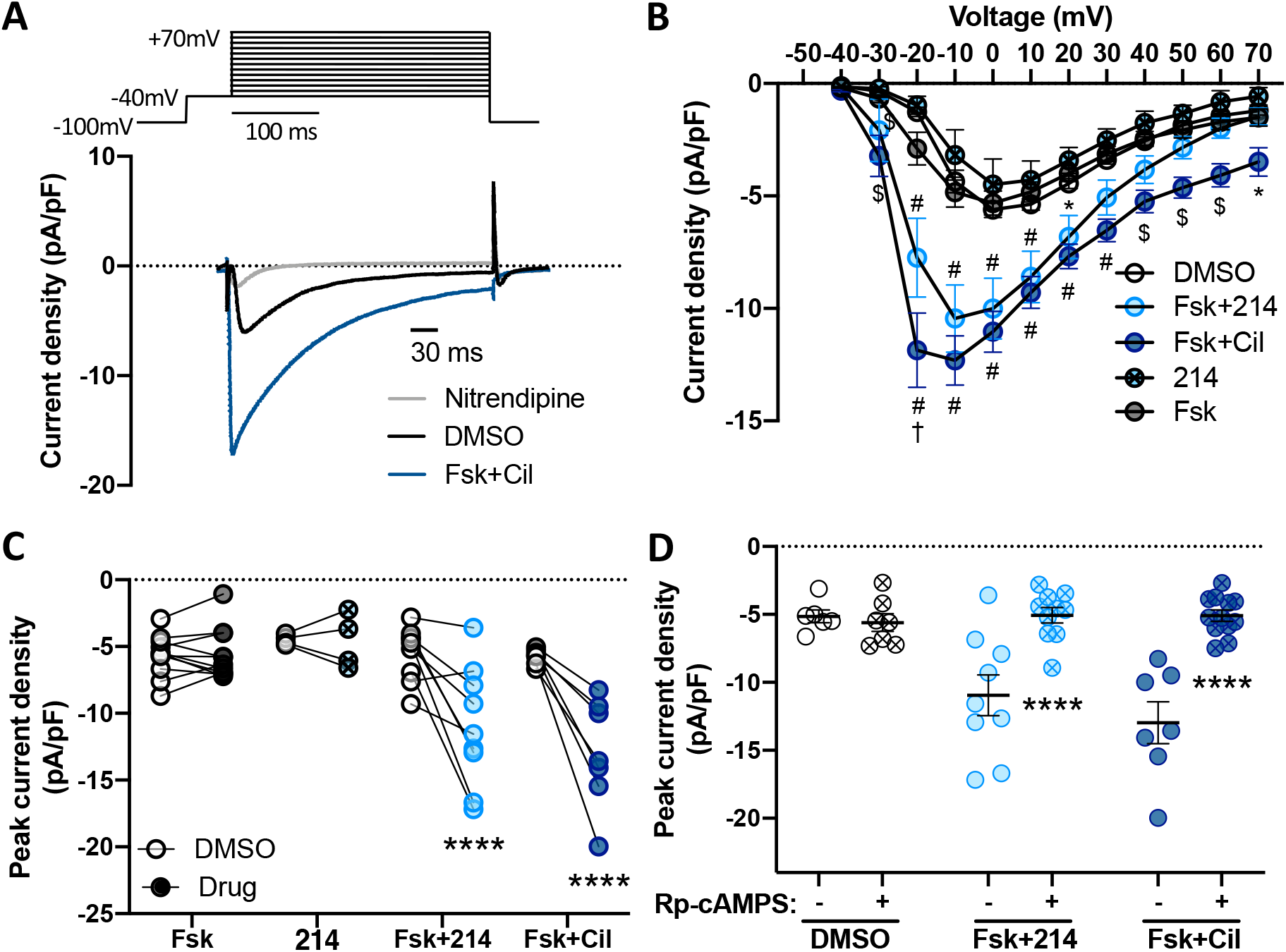
The L-type Ca^2+^ channel (LTCC) current increases upon PDE1 or 3 inhibition. LTCC current was measured in guinea pig myocytes. Cells were stimulated with the indicated drugs, and the inward current measured using the whole-cell voltage clamp protocol. Nitrendipine (10 μM) was used to confirm LTCC as the primary source of the current. A) The voltage-clamping protocol, and a representative trace showing change upon Fsk+Cil treatment, followed by sensitivity to nitrendipine are shown. B) The averaged peak current density values are plotted over a range of membrane voltage. *p<0.05, $p<0.001, #p<0.0001 against DMSO; †p<0.0001 against Fsk+214, ordinary 2-way ANOVA with Tukey’s multiple test. C) Paired-response plotted for change in the peak current density for cells before and after the indicated treatment. ****p<0.0001 vs respective DMSO, RM 2-way ANOVA with Sidak’s multiple test. D) Cells were again stimulated with indicated drugs 8-10 minutes after dialysis with Rp-cAMPS (100 μM). Data points from C are plotted again for comparison without PKA inhibition for change in the peak current density. ****p<0.0001 vs respective drug treatment without Rp-cAMPS; ordinary 2-way ANOVA with Sidak’s multiple test.

Given Fsk+PDE1i augmented Ca_v_1.2 current similarly as Fsk+PDE3i but without the latter’s engagement of the SR, we postulated that Fsk+PDE1i would be more sensitive to nitrendipine. Doses of nitrendipine that depressed but did not prevent myocyte shortening were determined in a range of 0.01–3 μM (Supplemental Fig 3A). Cells were treated with either DMSO, Fsk+PDE1i, or PDE3i (Fig 6A and B) with increasing nitrendipine dose. Fsk+PDE1/3i-stimulated shortening and peak Ca^2+^ transients with either inhibitor exhibited a nitrendipine dose-dependent decline. However, this sensitivity was greater in cells stimulated with Fsk+PDE1i as determined by a significantly steeper dose-response (Fig 6C). This was statistically significant as determined in log-transformed relations by analysis of covariance (Fig 6C and D).

**Fig 6.**
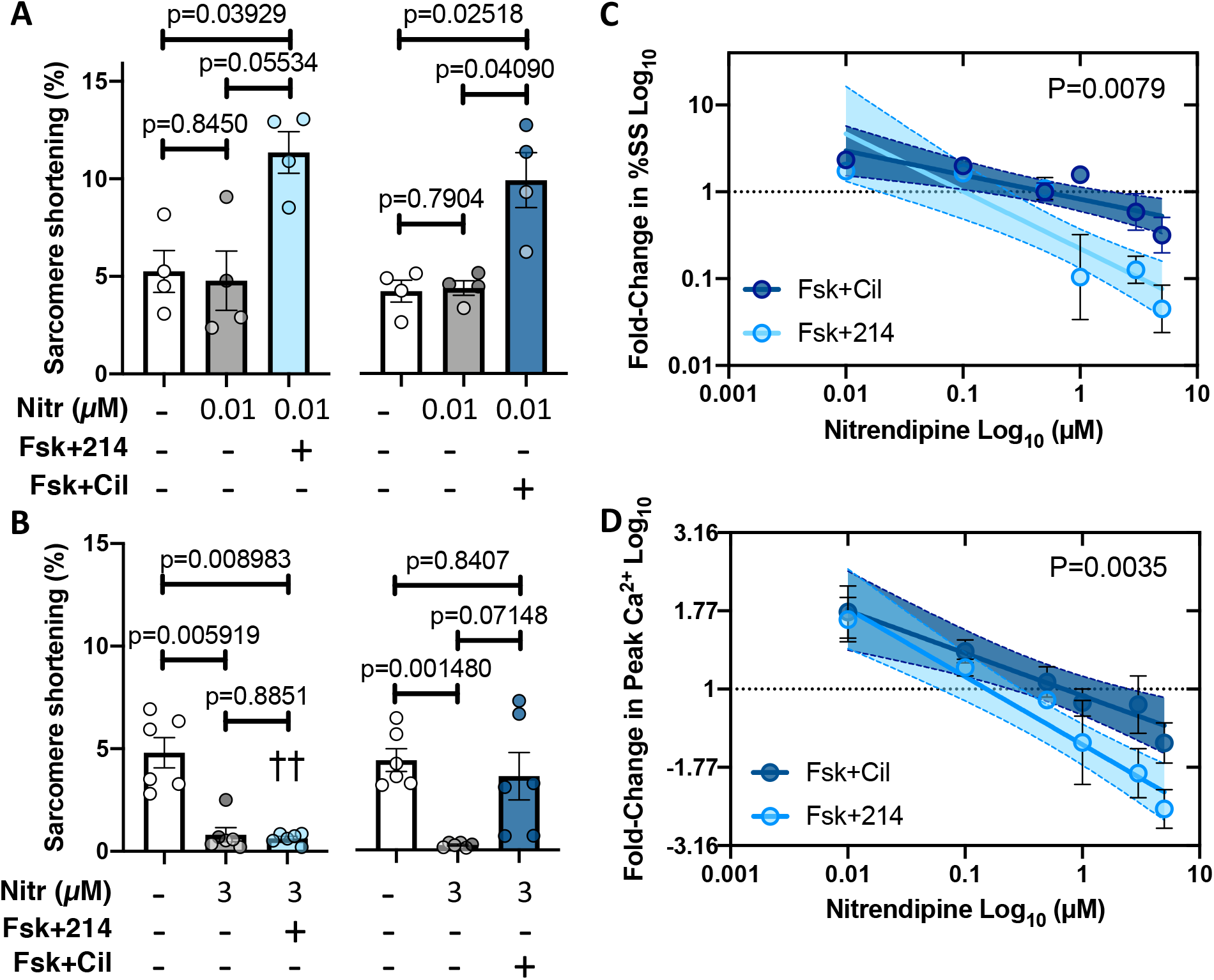
The inotropic effects of PDE1 is more sensitive to the LTCC blocker nitrendipine compared to that of PDE3. Cells were pre-treated with nitrendipine (nitr), before being stimulated further with Fsk+214 or Fsk+Cil. Change in sarcomere shortening response at nitr dose of A) 0.01 μM or B) 3 μM are plotted. The fold-difference in C) sarcomere shortening or D) peak Ca^2+^ transient (e.g., comparing nitr+Fsk+214 or nitr+Fsk+Cil vs DMSO) is plotted against Nitr dose, both expressed on logarithmic scale. The linear fit and 95% CI values are shown. P value in C indicates the difference in the slope, and that in D indicates the difference in the elevation or intercept; simple linear regression.

### PDE1i is less arrhythmogenic than PDE3i

Increased SR Ca^2+^ load and/or release is viewed as a potential cause for arrhythmia and attributed to pro-arrhythmia with both β-AR and PDE3i. As PDE1i did not alter SR Ca^2+^ load (unlike PDE3i), we hypothesized it may also be less arrhythmogenic despite increasing Ca_v_1.2 Ca^2+^ conductance. Myocytes treated with PDE1i or PDE3i in the presence of 0, 10, or 100 nM Fsk were examined for rhythm stability (fixed amplitude and rate under paced conditions) or arrhythmia (both parameters showing variability despite pacing). Figure 7A shows example tracings, and Figure 7B shows summary data for the relative percent of stable versus arrhythmic cells. PDE3i elicited more arrhythmia at lower Fsk doses than did PDE1i. This was most apparent without Fsk (81% vs 5%, p<0.0001) but persisted despite Fsk co-stimulation.

**Fig 7.**
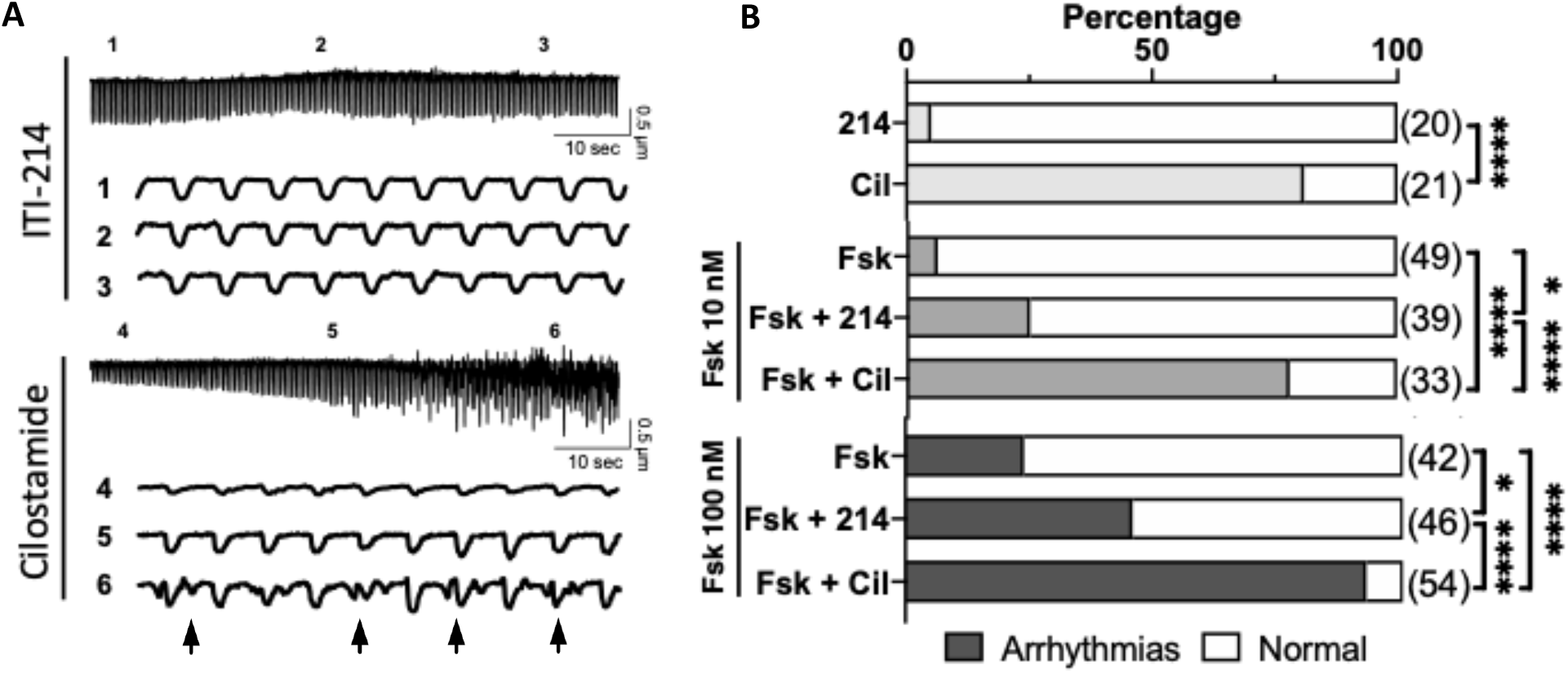
PDE1-I is less arrhythmogenic in comparison to PDE3-I. Guinea pig myocytes treated with 214 or Cil in the presence of Fsk were scored as normal or arrhythmic. A) Representative sarcomere shortening trace for cells responding to 214 (top) or Cil (bottom). Numbers 1-6 indicate regions of interest, with sarcomere shortening traces corresponding to each region shown underneath. Arrows indicate delayed afterdepolarizations (DADs) in cells treated with Cil. B) The percentage of cells in arrhythmic contractions (shades) or normal (white) for indicated conditions.

## Discussion

This study reveals the mechanisms of inotropic modulation in cardiac myocytes from guinea pig that express PDE1C by a broad PDE1i and contrasts the findings to those from PDE3i. As summarized in Figure 8, we find PDE1i augments contractility in a PKA dependent manner by increasing Ca_v_1.2 current density; however, in contrast to β-ARs, PDE1i does not desensitize the myofilaments to Ca^2+^. Thus, while we find no increase in SR Ca^2+^ load or PLN phosphorylation, contraction still increases with less total intracellular Ca^2+^ rise as compared to β-ARs or PDE3i. This profile is further supported by greater sensitivity of PDE1i positive inotropy to Ca_v_1.2 blockade. An important correlate is less arrhythmogenicity than observed with PDE3i. These findings deepen our understanding of PDE1 regulation in myocytes and have translational significance as the first clinical trial of a PDE1i, ITI-214, in heart failure patients has found it safe and to enhance contraction ^10^.

**Fig 8.**
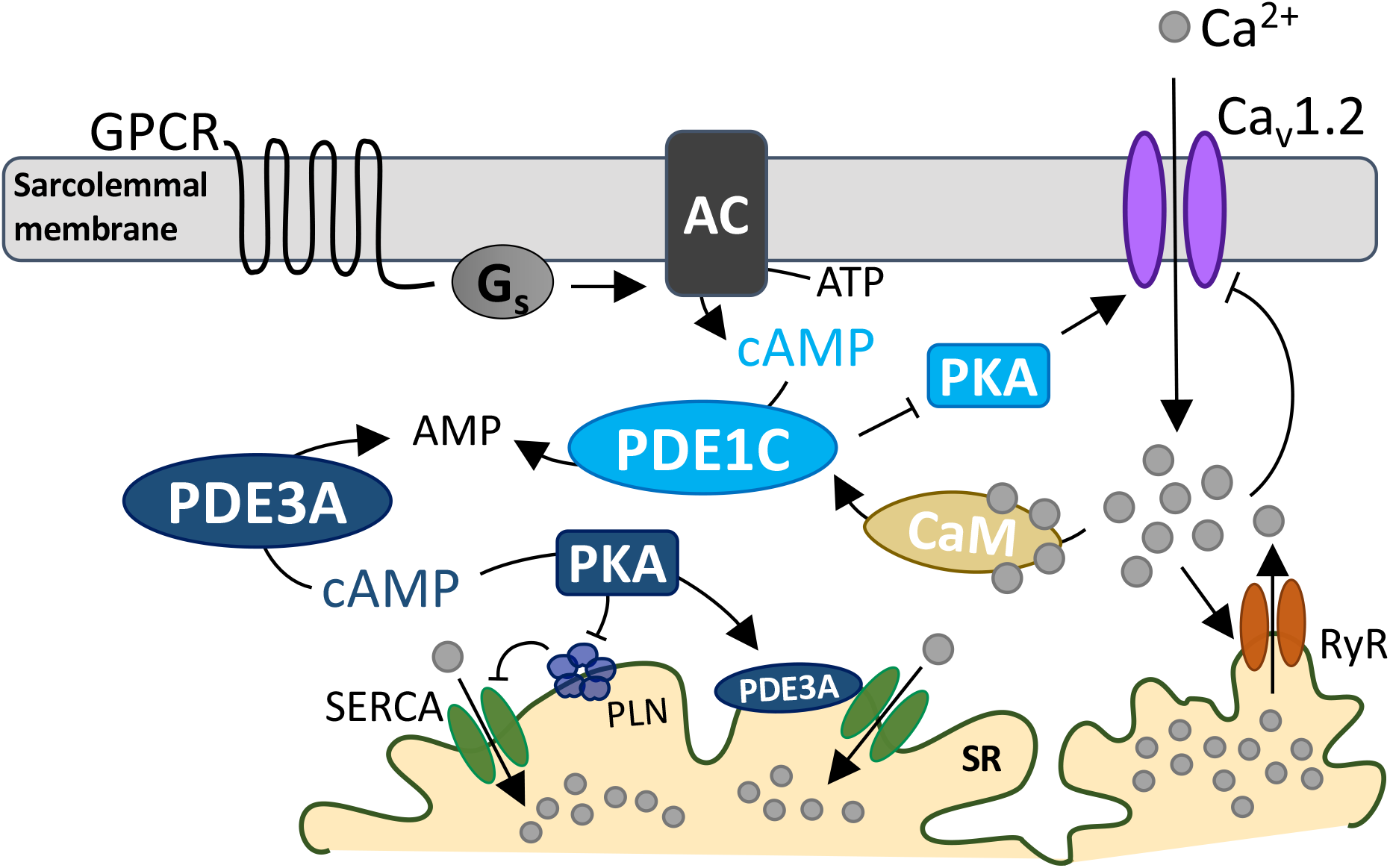
Schematic of the proposed working model. PDE1C may be part of a negative feedback mechanism controlling Ca^2+^ influx via Ca_v_1.2. Downstream of Ca_v_1.2 activation by a GPCR, PDE1C senses an increase in LTCC activity putatively via Ca^2+^/CaM. PDE1C hydrolyzes cAMP to AMP to relieve the stimulatory effects of PKA upon Ca_v_1.2activity. This negative feedback mechanism may operate in conjunction with Ca^2+^-induced Ca_v_1.2 inactivation. In contrast, PDE3A hydrolyzes a different pool of cAMP at the SR to increase PKA phosphorylation of PLN to release its inhibition of SERCA-mediated Ca^2+^ reuptake. This likely occurs in conjunction with PKA phosphorylation of PDE3A to directly increase SERCA activity. Abbreviations: G-protein coupled receptor (GPCR), stimulatory G protein (Gs), adenylyl cyclase (AC), phosphodiesterase 1C (PDE1C), phosphodiesterase 3A (PDE3A), protein kinase A (PKA), calmodulin (CaM), sarcoplasmic reticulum (SR), SR Ca2+ ATPase (SERCA), phospholamban (PLN), ryanodine receptor (RyR).

### PDE modulators of myocyte contractility

Acute augmentation of cardiac contractility is primarily coupled to pathways that activate cAMP-PKA signaling. This is intrinsic to β-AR stimulation and other G-protein coupled receptors such as sub-types of adenosine receptors and glucagon receptors ^34^. As cAMP itself is the same molecule throughout the cell its signaling requires exquisite local control. This is provided in part by specific cAMP-hydrolyzing PDEs ^35^. In larger mammalian hearts, PDE3 plays a dominant role, whereas PDE4 is more influential in mouse and rat. Until recently, the role of PDE1 in larger mammalian myocytes and hearts was unknown. PDE3 binds cAMP in the nM range ^36^ whereas PDEs 1 and 4 operate at μM ranges ^9, 37^. As intracellular cAMP concentration is ~ 1 μM ^38^, PDE3 inhibitors are expected to have more impact under resting conditions even in isolated myocytes. By contrast, although PDE1i *in vivo* augments contractility, in isolated cells devoid of adrenergic tone, cAMP must first be elevated to observe inotropy as show here in guinea pig and previously in mouse and rabbit ^8, 39^.

### Evidence for distinct PDE1 vs PDE3 localization

Beyond their relative affinity for cAMP, differential regulation by spatially compartmentalized PDEs control local cAMP-PKA signaling ^40^. PDE3 is found at the plasma membrane ^41^ and also at the SR where it controls local Ca^2+^ uptake ^15^. PDE1 also localizes to the plasma membrane and displays immunofluorescence staining along Z- and M-lines in human myocytes ^42^. However, PDE1 accounts for only 14% of microsomal PDE activity against cAMP as compared to 78% residing in the cytoplasmic fraction. This contrasts with PDE3, which contributes 69% of cAMP esterase activity in large mammal cardiac microsomal fractions that include the SR ^42^. These differences could underlie the lack of PLN phosphorylation and associated SR Ca^2+^ content modulation by PDE1i observed here.

PDE3i or PDE1i increased Ca_v_1.2 current similarly, placing both PDEs within caveolin-enriched microdomains of the sarcolemma, ^43, 44^ where this channel resides ^45^. However, the two PDEs displayed marked differences in their interaction with β-AR agonism, with PDE3i augmenting both Ca^2+^ and shortening when combined with Iso but PDE1i having little impact. This suggests that while proximate to the channel, the two PDEs reside in different nanodomains relative to G_s_-coupled GPCRs. Beta-adrenergic stimulated Ca_v_1.2 current requires Rad phosphorylation which otherwise constitutively suppresses current via a complex with the channel’s β and α_1C_ subunits ^11, 12^. Proximity protein analysis has identified PDE4 isoforms, PDE3A, and PDE1C as being near Ca_v_1.2. PDE1C is also found in complex with adenosine A_2_ receptors and the non-selective cation channel, transient receptor potential cation channel 3 (TRPC3) where it plays a role in cytoprotection signaling^46^.

### PDE1 and PDE3 differentially impact intracellular PKA signaling

While our results indicate that β-AR G_s_-coupled cAMP synthesis is regulated by PDE3 rather than PDE1, the sources of cAMP that are more selectively controlled by PDE1 remain to be fully elucidated. As noted, one source may be A_2_ adenosine receptors, as A_2B_R blockade *in vivo* prevented positive inotropy in the rabbit ^8^, and PDE1C modulation of A_2_R signaling impacts cAMP-dependent cardioprotection against doxorubicin-mediated cell death ^46^. In the latter case, the Ca^2+^ trigger was coupled to transient receptor potential canonical channel type 3 (TRPC3) identified in a protein complex with PDE1C and A_2_R. Interestingly, TRPC3 modulation of myocyte hypertrophy is linked to activation of nuclear factor of activated T-cell (NFAT) that has been shown to require Ca_v_1.2 current as it is suppressed by nifedipine ^47^. This crosstalk works in both directions, as genetic upregulation of TRPC3 slows the decay of the Ca_v_1.2 current, augmenting the net Ca^2+^ transient. The present data showing PDE1 modulation of this channel along with data linking it to TRPC3 suggest that this localized complex likely impacts contractility as well as hypertrophic remodeling.

PDE3i increased arrhythmogenicity more than PDE1i, and importantly this disparity persisted even despite higher levels of basal cAMP stimulation. This resonates with clinical findings, as prior heart failure trials with PDE3i identified malignant arrhythmia as a significant risk ^7^. Sudden cardiac death and increased ventricular ectopy were observed in canine and human HF patients receiving milrinone ^48–50^. In the recently completed clinical trial testing ITI-214 efficacy and tolerance in heart failure patients (NCT03387215), no change in arrhythmias were noted ^10^. These findings are in agreement with results in both rabbit and dog *in vivo*, where pro-arrhythmia was not detected ^8^, while both studies observed increased contractility and vascular dilation with ITI-214 treatment.

### Limitations

While this study reveals novel mechanisms for PDE1i inotropy, a number of questions remain to be addressed. The precise localization of the PDE1 regulated cAMP nanodomains, and identity of the proteins directly impacted by its inhibition are unknown. Studies employing locally targeted fluorescent resonance energy transfer probes and proximity labeling and phospho-proteomics aim to address these unknowns. Calcium homeostasis is tightly controlled, and while we have shown PDE1i increases Ca_v_1.2 current, this must be balanced by removal via either the SR, the Na^+^/Ca^2+^ exchanger, or mitochondrial Ca^2+^ uniporter^32^. Whether PDE1i enhances either of the latter two removal mechanisms remains untested, though unlike the SR, they are not thought to be significantly altered by PKA activation^51^ as required for the PDE1i effect. It remains possible that some enhanced SR uptake occurs that is below the detection level of the caffeine release method used here, and we recognize that beyond the patch-clamp studies, direct evidence of a local pool of inward Ca^2+^ with PDE1i in beating cells remains to be obtained. Nanodomain targeted Ca^2+^ sensor studies might address this in future studies. Lastly, while the contraction/calcium responses were PKA dependent and speak to cAMP as the primary modulated species, some role for cGMP remains. PDE1A, which preferentially hydrolyzes cGMP >20x more than cAMP, is also expressed in larger mammals and humans. While very potent, ITI-214 is not PDE1 isoform selective ^52^ since the catalytic site is highly homologous among the isoforms. However, cGMP elevation has not been demonstrated to augment Ca_v_1.2, if anything the opposite effect has been reported ^53^.

## Conclusions

In summary, we show that similar to PDE3i, PDE1i increases Ca_v_1.2 activity in a PKA-dependent manner, increasing myocyte contraction in cells exposed to background adenylyl cyclase stimulation though not that coupled to β-AR. Unlike PDE3i or β-AR stimulation, PLN, TnI, and MyBP-C are not phosphorylated upon PDE1i. Our findings support PDE1 as a regulator of Ca_v_1.2 that is likely leveraged by the lack of concomitant myofilament Ca^2+^ desensitization, resulting in cAMP/PKA dependent inotropy that is less arrhythmogenic. Ongoing studies at basic and clinical levels will continue to test the potential of this intervention as a heart failure therapy and further clarify its signaling components.

### ACKNOWLEDGEMENTS

The authors would like to thank: Dr. Peng Li for providing ITI-214 used in these studies; Drs. Ting Liu and Brian O’Rourke for sharing their expertise in guinea pig handling and myocyte isolation techniques; Dr. Sakthivel Sadayappan for his generous gift of myosin binding protein-C antibodies; Drs. Kobra Haghighi and Litsa Kranias for technical advice on phospholamban western blotting; and Ms. Laurel Keefer for her assistance in R coding language.

### SOURCES OF FUNDING

This work was supported by research support from Intracellular Therapies, Inc., Grants AHA 18POST33960157 and 20CDA35260135 to GKM, R35 HL140034 to MEA, and AHA AWRP 17CSA33610107 and NIH/NHLBI R35 HL140034 to DAK.

### DISCLOSURES

David Kass is a consultant for Intracellular Therapies, Inc.

### Supplemental Materials

Online Figures 1 - 3

## Non-standard Abbreviations and Acronyms

β-ARs: (β adrenergic receptor stimulation)
PDE1i: (phosphodiesterase type 1 inhibition)
PDE3i: (phosphodiesterase type 3 inhibition)
PLN: phospholamban
SR: sarcoplasmic reticulum
TnI: troponin I
MyBPC: myosin binding protein C

## Notes

### Competing Interest Statement

Dr. David Kass is a consultant with Intracellular Therapies Inc., manufacturer of ITI-214 that is used in this study.

## REFERENCES

1. Conrad N, Judge A, Tran J, Mohseni H, Hedgecott D, Crespillo AP, Allison M, Hemingway H, Cleland JG, McMurray JJV and Rahimi K. Temporal trends and patterns in heart failure incidence: a population-based study of 4 million individuals. Lancet. 2018;391:572–580.

2. Malik FI, Hartman JJ, Elias KA, Morgan BP, Rodriguez H, Brejc K, Anderson RL, Sueoka SH, Lee KH, Finer JT, Sakowicz R, Baliga R, Cox DR, Garard M, Godinez G, Kawas R, Kraynack E, Lenzi D, Lu PP, Muci A, Niu C, Qian X, Pierce DW, Pokrovskii M, Suehiro I, Sylvester S, Tochimoto T, Valdez C, Wang W, Katori T, Kass DA, Shen YT, Vatner SF and Morgans DJ. Cardiac myosin activation: a potential therapeutic approach for systolic heart failure. Science. 2011;331:1439–43.

3. Movsesian MA and Bristow MR. Alterations in cAMP-mediated signaling and their role in the pathophysiology of dilated cardiomyopathy. CurrTopDevBiol. 2005;68:25–48.

4. Bistola V, Arfaras-Melainis A, Polyzogopoulou E, Ikonomidis I and Parissis J. Inotropes in Acute Heart Failure: From Guidelines to Practical Use: Therapeutic Options and Clinical Practice. Card Fail Rev. 2019;5:133–139.

5. Felker GM, Benza RL, Chandler AB, Leimberger JD, Cuffe MS, Califf RM, Gheorghiade M, O’Connor CM and Investigators O-C. Heart failure etiology and response to milrinone in decompensated heart failure: results from the OPTIME-CHF study. J Am Coll Cardiol. 2003;41:997–1003.

6. Abraham W, Adams K, Fonarow G, Costanzo M, Berkowitz R, LeJemtel T, Cheng M, Wynne J, Investigators ASACa and Group AS. In-hospital Mortality in Patients With Acute Decompensated Heart Failure Requiring Intravenous Vasoactive Medications: An Analysis From the Acute Decompensated Heart Failure National Registry (ADHERE). Journal of the American College of Cardiology. 2005;46:57–64.

7. Packer M, Carver JR, Rodeheffer RJ, Ivanhoe RJ, DiBianco R, Zeldis SM, Hendrix GH, Bommer WJ, Elkayam U, Kukin ML and et al. Effect of oral milrinone on mortality in severe chronic heart failure. The PROMISE Study Research Group. N Engl J Med. 1991;325:1468–75.

8. Hashimoto T, Kim GE, Tunin RS, Adesiyun T, Hsu S, Nakagawa R, Zhu G, O’Brien JJ, Hendrick JP, Davis RE, Yao W, Beard D, Hoxie HR, Wennogle LP, Lee DI and Kass DA. Acute Enhancement of Cardiac Function by Phosphodiesterase Type 1 Inhibition. Circulation. 2018;138:1974–1987.

9. Bender AT and Beavo JA. Cyclic nucleotide phosphodiesterases: molecular regulation to clinical use. Pharmacol Rev. 2006;58:488–520.

10. Gilotra NA, Devore A, Hays A, Hahn V, Agunbiade TA, Chen R, Davis R, Satlin A, Povsic T and Kass D. Cardiac and Hemodynamic Effects of Acute Phosphodiesterase-1 Inhibition in Human Heart Failure. Journal of Cardiac Failure. 2020;26:S12.

11. Liu G, Papa A, Katchman A, Zakharov S, Roybal D, Hennessey J, Kushner J, Yang L, Chen B, Kushnir A, Dangas K, Gygi S, Pitt G, Colecraft H, Ben-Johny M, Kalocsay M and Marx S. Mechanism of Adrenergic Ca V 1.2 Stimulation Revealed by Proximity Proteomics. Nature. 2020;577:695–700.

12. Papa A, Kushner JS, Hennessey JA, Katchman AN, Zakharov SI, Chen B-X, Yang L, Lu R, Leong S, Diaz J, Liu G, Roybal DD, Liao X, del Rivero Morfin P, Colecraft HM, Pitt GS, Clarke OB, V.K. T, Johny MB and Marx SO. Adrenergic Ca_V_1.2 Activation via Rad Phosphorylation Converges at alpha_1C_ I-II Loop. Circulation Research. 2020;0.

13. Beca S, Ahmad F, Shen W, Liu J, Makary S, Polidovitch N, Sun J, Hockman S, Chung YW, Movsesian M, Murphy E, Manganiello V and Backx PH. Phosphodiesterase type 3A regulates basal myocardial contractility through interacting with sarcoplasmic reticulum calcium ATPase type 2a signaling complexes in mouse heart. Circ Res. 2013;112:289–97.

14. Haghighi K, Kolokathis F, Gramolini AO, Waggoner JR, Pater L, Lynch RA, Fan GC, Tsiapras D, Parekh RR, Dorn GW, 2nd, MacLennan DH, Kremastinos DT and Kranias EG. A mutation in the human phospholamban gene, deleting arginine 14, results in lethal, hereditary cardiomyopathy. Proc Natl Acad Sci U S A. 2006;103:1388–93.

15. Ahmad F, Shen W, Vandeput F, Szabo-Fresnais N, Krall J, Degerman E, Goetz F, Klussmann E, Movsesian M and Manganiello V. Regulation of sarcoplasmic reticulum Ca2+ ATPase 2 (SERCA2) activity by phosphodiesterase 3A (PDE3A) in human myocardium: phosphorylation-dependent interaction of PDE3A1 with SERCA2. J Biol Chem. 2015;290:6763–76.

16. Layland J, Solaro RJ and Shah AM. Regulation of cardiac contractile function by troponin I phosphorylation. Cardiovasc Res. 2005;66:12–21.

17. Tong CW, Stelzer JE, Greaser ML, Powers PA and Moss RL. Acceleration of crossbridge kinetics by protein kinase A phosphorylation of cardiac myosin binding protein C modulates cardiac function. CircRes. 2008;103:974–982.

18. Nagayama T, Takimoto E, Sadayappan S, Mudd JO, Seidman JG, Robbins J and Kass DA. Control of in vivo left ventricular contraction/relaxation kinetics by myosin binding protein C: protein kinase A phosphorylation dependent and independent regulation. Circulation. 2007;116:2399–2408.

19. Liu T, Takimoto E, Dimaano VL, DeMazumder D, Kettlewell S, Smith G, Sidor A, Abraham TP and O’Rourke B. Inhibiting Mitochondrial Na+/Ca2+ Exchange Prevents Sudden Death in a Guinea Pig Model of Heart Failure. Circ Res. 2014;115:44–54.

20. Isenberg G and Klockner U. Calcium tolerant ventricular myocytes prepared by preincubation in a “KB medium”. Pflugers Arch. 1982;395:6–18.

21. Ellingsen O, Davidoff AJ, Prasad SK, Berger HJ, Springhorn JP, Marsh JD, Kelly RA and Smith TW. Adult rat ventricular myocytes cultured in defined medium: phenotype and electromechanical function. Am J Physiol. 1993;265:H747–54.

22. Money-Kyrle AR, Davies CH, Ranu HK, O’Gara P, Kent NS, Poole-Wilson PA and Harding SE. The role of cAMP in the frequency-dependent changes in contraction of guinea-pig cardiomyocytes. Cardiovasc Res. 1998;37:532–40.

23. Tang M, Zhang X, Li Y, Guan Y, Ai X, Szeto C, Nakayama H, Zhang H, Ge S, Molkentin JD, Houser SR and Chen X. Enhanced basal contractility but reduced excitation-contraction coupling efficiency and beta-adrenergic reserve of hearts with increased Cav1.2 activity. Am J Physiol Heart Circ Physiol. 2010;299:H519–28.

24. Kirk JA, Holewinski RJ, Kooij V, Agnetti G, Tunin RS, Witayavanitkul N, de Tombe PP, Gao WD, Van Eyk J and Kass DA. Cardiac resynchronization sensitizes the sarcomere to calcium by reactivating GSK-3beta. J Clin Invest. 2014;124:129–38.

25. Hsu S, Kokkonen-Simon KM, Kirk JA, Kolb TM, Damico RL, Mathai SC, Mukherjee M, Shah AA, Wigley FM, Margulies KB, Hassoun PM, Halushka MK, Tedford RJ and Kass DA. Right Ventricular Myofilament Functional Differences in Humans With Systemic Sclerosis-Associated Versus Idiopathic Pulmonary Arterial Hypertension. Circulation. 2018;137:2360–2370.

26. Rasmussen T, Wu Y, Joiner M, Koval O, Wilson N, Luczak E, Wang Q, Chen B, Gao Z, Zhu Z, Wagner B, Soto J, McCormick M, Kutschke W, Weiss R, Yu L, Boudreau R, Abel E, Zhan F, Spitz D, Buettner G, Song L, Zingman L and Anderson M. Inhibition of MCU Forces Extramitochondrial Adaptations Governing Physiological and Pathological Stress Responses in Heart. Proceedings of the National Academy of Sciences of the United States of America. 2015;112:9129–9134.

27. Richter W, Xie M, Scheitrum C, Krall J, Movsesian M and Conti M. Conserved Expression and Functions of PDE4 in Rodent and Human Heart. Basic research in cardiology. 2011;106:249–262.

28. Sprenger JU, Bork NI, Herting J, Fischer TH and Nikolaev VO. Interactions of Calcium Fluctuations during Cardiomyocyte Contraction with Real-Time cAMP Dynamics Detected by FRET. PLoS One. 2016;11:e0167974.

29. Gambassi G, Spurgeon H, Lakatta E, Blank P and Capogrossi M. Different Effects of Alpha- And Beta-Adrenergic Stimulation on Cytosolic pH and Myofilament Responsiveness to Ca2+ in Cardiac Myocytes. Circulation research. 1992;71:870–882.

30. Mohamed AS, Dignam JD and Schlender KK. Cardiac myosin-binding protein C (MyBP-C): identification of protein kinase A and protein kinase C phosphorylation sites. Arch Biochem Biophys. 1998;358:313–9.

31. Mackiewicz UL, B. Temperature dependent contribution of Ca2+ transporters to relaxation in cardiac myocytes: important role of sarcolemmal Ca2+-ATPase. J Physiol Pharmacol 2006;Mar;57:3–15.

32. Bers DM. Calcium cycling and signaling in cardiac myocytes. AnnuRevPhysiol. 2008;70:23–49.

33. Pogwizd SM, Schlotthauer K, Li L, Yuan W and Bers DM. Arrhythmogenesis and contractile dysfunction in heart failure: Roles of sodium-calcium exchange, inward rectifier potassium current, and residual beta-adrenergic responsiveness. Circ Res. 2001;88:1159–67.

34. Pinilla-Vera M, Hahn VS and Kass DA. Leveraging Signaling Pathways to Treat Heart Failure With Reduced Ejection Fraction. Circ Res. 2019;124:1618–1632.

35. Kim GE and Kass DA. Cardiac Phosphodiesterases and Their Modulation for Treating Heart Disease. In: J. Bauersachs, J. Butler and P. Sandner, eds. Heart Failure. 2016/10/28 ed.: Springer, Cham; 2017(243): 249–269.

36. Manganiello V, Murata T, Taira M, Belfrage P and Degerman E. Diversity in Cyclic Nucleotide Phosphodiesterase Isoenzyme Families. Archives of biochemistry and biophysics. 1995;322:1–13.

37. Salanova M, Jin S and Conti M. Heterologous Expression and Purification of Recombinant Rolipram-Sensitive Cyclic AMP-specific Phosphodiesterases. Methods (San Diego, Calif). 1998;14:55–64.

38. Koschinski A and Zaccolo M. Quantification and Comparison of Signals Generated by Different FRET-Based cAMP Reporters. Methods Mol Biol. 2019;1947:217–237.

39. Sprenger JU, Bork NI, Herting J, Fischer TH and Nikolaev VO. Interactions of Calcium Fluctuations during Cardiomyocyte Contraction with Real-Time cAMP Dynamics Detected by FRET. PLOS ONE. 2016;11.

40. Surdo NC, Berrera M, Koschinski A, Brescia M, Machado MR, Carr C, Wright P, Gorelik J, Morotti S, Grandi E, Bers DM, Pantano S and Zaccolo M. FRET biosensor uncovers cAMP nano-domains at β-adrenergic targets that dictate precise tuning of cardiac contractility. Nature Communications. 2017;8:15031.

41. Sun B, Li H, Shakur Y, Hensley J, Hockman S, Kambayashi J, Manganiello VC and Liu Y. Role of phosphodiesterase type 3A and 3B in regulating platelet and cardiac function using subtype-selective knockout mice. Cell Signal. 2007;19:1765–71.

42. Vandeput F, Wolda SL, Krall J, Hambleton R, Uher L, McCaw KN, Radwanski PB, Florio V and Movsesian MA. Cyclic nucleotide phosphodiesterase PDE1C1 in human cardiac myocytes. J Biol Chem. 2007;282:32749–57.

43. Kawai M, Hussain M and Orchard C. Excitation-contraction Coupling in Rat Ventricular Myocytes After Formamide-Induced Detubulation. The American journal of physiology. 1999;277:H603–609.

44. Pásek M, Brette F, Nelson A, Pearce C, Qaiser A, Christe G and Orchard C. Quantification of T-Tubule Area and Protein Distribution in Rat Cardiac Ventricular Myocytes. Progress in biophysics and molecular biology. 2008;96:244–257.

45. Bhargava A, Lin X, Novak P, Mehta K, Korchev Y, Delmar M and Gorelik J. Super-resolution scanning patch clamp reveals clustering of functional ion channels in adult ventricular myocyte. Circ Res. 2013;112:1112–1120.

46. Zhang Y, Knight W, Chen S, Mohan A and Yan C. A Multi-Protein Complex with TRPC, PDE1C, and A2R Plays a Critical Role in Regulating Cardiomyocyte cAMP and Survival. Circulation. 2018.

47. Gao H, Wang F, Wang W, Makarewich CA, Zhang H, Kubo H, Berretta RM, Barr LA, Molkentin JD and Houser SR. Ca(2+) influx through L-type Ca(2+) channels and transient receptor potential channels activates pathological hypertrophy signaling. J Mol Cell Cardiol. 2012;53:657–67.

48. Kittleson M, Johnson L and Pion P. The Acute Hemodynamic Effects of Milrinone in Dogs With Severe Idiopathic Myocardial Failure. Journal of veterinary internal medicine. 1987;1:121–127.

49. Holmes J, Kubo S, Cody R and Kligfield P. Milrinone in Congestive Heart Failure: Observations on Ambulatory Ventricular Arrhythmias. American heart journal. 1985;110:800–806.

50. Anderson J, Askins J, Gilbert E, Menlove R and Lutz J. Occurrence of Ventricular Arrhythmias in Patients Receiving Acute and Chronic Infusions of Milrinone. American heart journal. 1986;111:466–474.

51. Zhang YH and Hancox JC. Regulation of cardiac Na+-Ca2+ exchanger activity by protein kinase phosphorylation--still a paradox? Cell Calcium. 2009;45:1–10.

52. Snyder GL, Prickaerts J, Wadenberg ML, Zhang L, Zheng H, Yao W, Akkerman S, Zhu H, Hendrick JP, Vanover KE, Davis R, Li P, Mates S and Wennogle LP. Preclinical profile of ITI-214, an inhibitor of phosphodiesterase 1, for enhancement of memory performance in rats. Psychopharmacology (Berl). 2016;233:3113–24.

53. Yang L, Liu G, Zakharov SI, Bellinger AM, Mongillo M and Marx SO. Protein kinase G phosphorylates Cav1.2 alpha1c and beta2 subunits. Circ Res. 2007;101:465–474.

