## Supplemental Figures for "PDE1 inhibition modulates Ca_v_1.2 channel to stimulate cardiomyocyte contraction"

**Supplemental Material**
Supplemental Fig 1-3

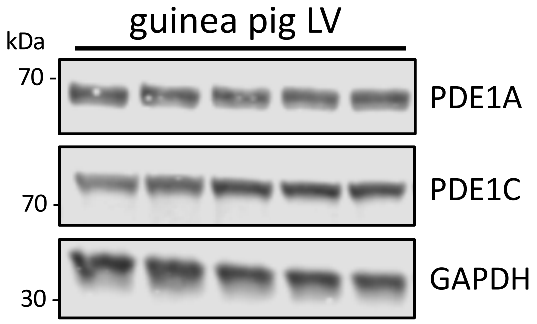


Supplemental Fig 1. **PDE1A and 1C expression in guinea pig myocytes.** The left ventricular tissue was homogenized, before being probed for PDE1A and PDE1C expression levels. GAPDH is shown as loading control.

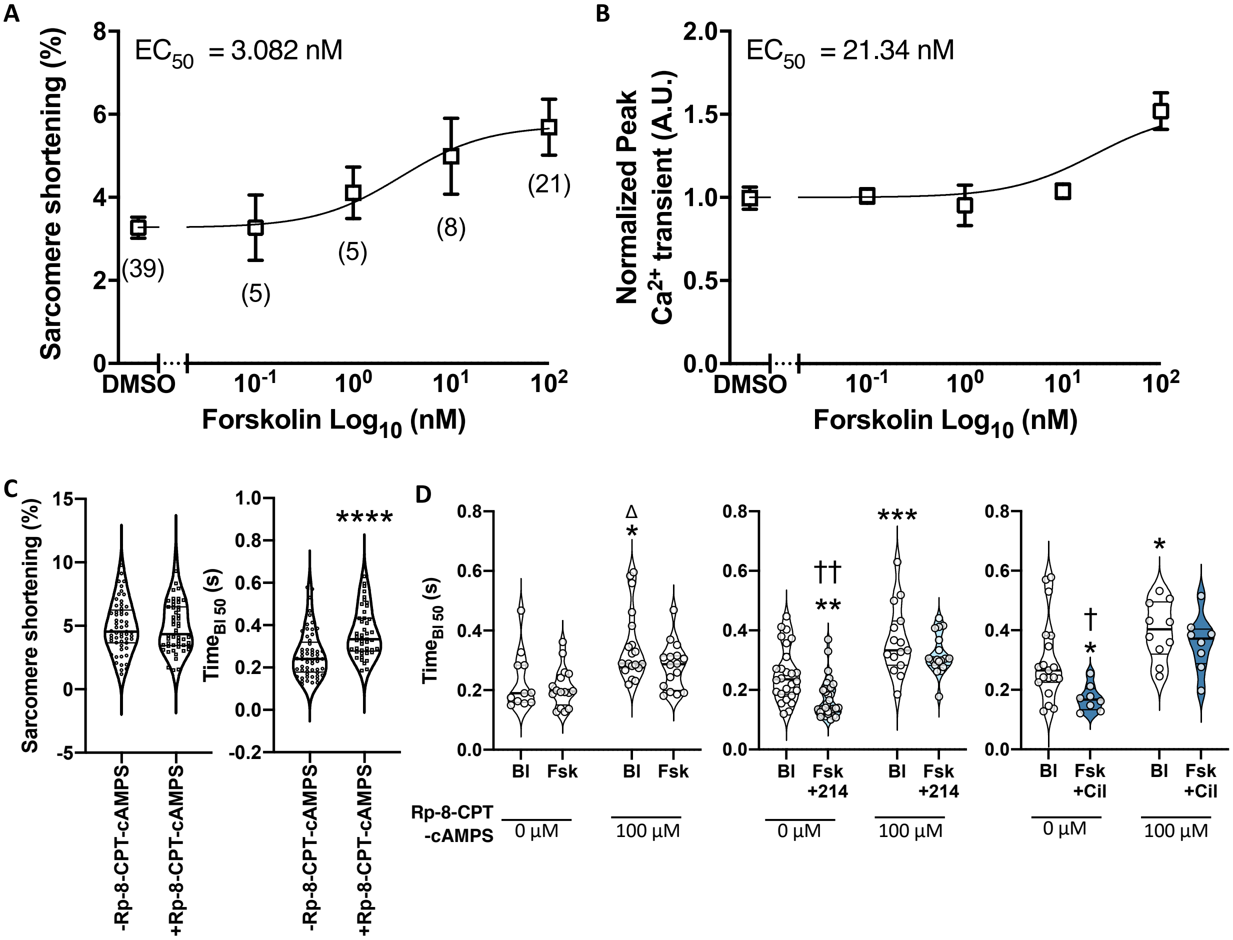


Supplemental Fig 2. **Dose response to forskolin**. Myocytes were treated with increasing amount of forskolin. Increase in A) the sarcomere shortening or B) peak $Ca^{2+}$ transient was plotted against forskolin dose, expressed on a logarithmic scale. The response was fitted with a nonlinear fit, and $EC_{50}$ values are reported. C) Baseline cell sarcomere shortening and relaxation kinetics (time to 50% of baseline) were compared in the absence or presence of 100 $\mu$M Rp-8-CPT-cAMPS; ****p<0.0001, unpaired 2-tailed t-test. D) Change in relaxation kinetics was compared in the absence or presence of 100 $\mu$M Rp-8-CPT-cAMPS for cells treated with Fsk (10 nM), Fsk (10 nM) + 214 (1 $\mu$M), or Fsk (10 nM) + Cil (1 $\mu$M). *p<0.05, **p<0.01, ***p<0.001 vs baseline (Bl) without Rp-8-CPT-cAMPS; Δp<0.001 vs Fsk without Rp-8-CPT-cAMPS; †p<0.01, ††p<0.0001 vs corresponding drug condition in the presence of Rp-8-CPT-cAMPS; ordinary 2-way ANOVA with Sidak's test.

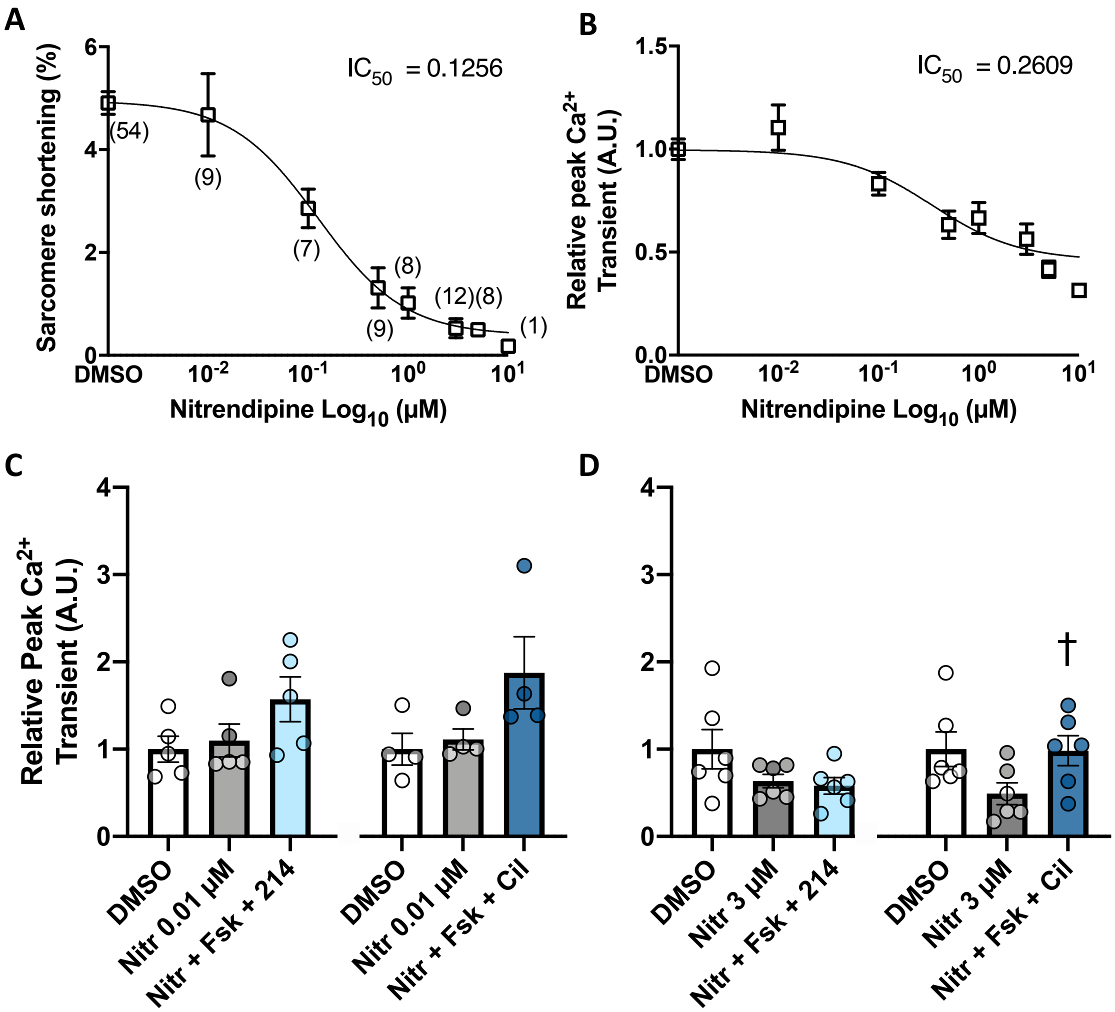


Supplemental Fig 3. **Dose response to nitrendipine**. Guinea pig myocytes were treated with increasing amount of nitrendipine (Nitr). Decreases in the A) sarcomere shortening or B) peak $Ca^{2+}$ transient values are plotted against nitrendipine dose, the latter of which is expressed on a logarithmic scale. The response was fitted with a nonlinear fit, and $EC_{50}$ values are reported. C and D) The peak $Ca^{2+}$ transient response is plotted for conditions in cells treated as indicated. †$p<0.05$ vs Nitr+Fsk+214, Uncorrected Fisher's LSD test.
